# Tobramycin-induced secretion of *P. aeruginosa* 5′ tRNA-fMet halves suppresses lung inflammation via AGO2 gene silencing

**DOI:** 10.1101/2021.09.23.461540

**Authors:** Zhongyou Li, Katja Koeppen, Alix Ashare, Deborah A. Hogan, Scott A. Gerber, Bruce A. Stanton

## Abstract

Although inhaled tobramycin increases lung function in people with cystic fibrosis (pwCF), the density of *P. aeruginosa* in the lungs is only modestly reduced by tobramycin; hence, the mechanism whereby tobramycin improves lung function is unclear. Here, we demonstrate that tobramycin increases the abundance of two 5′ tRNA-fMet halves in outer membrane vesicles (OMVs) secreted by *P. aeruginosa* and that the 5′ tRNA-fMet halves reduce IL-8 secretion by CF bronchial epithelial cells (CF-HBECs). In mouse lung, the 5′ tRNA-fMet halves attenuate KC secretion and neutrophil recruitment. We also report that the 5′ tRNA-fMet halves suppress pro-inflammatory network gene expression by an Argonaut 2 (AGO2)-mediated gene silencing mechanism, thereby reducing IL-8 secretion in CF-HBECs. Moreover, tobramycin reduces the IL-8 concentration and neutrophil content in bronchoalveolar lavage fluid of pwCF. Thus, we conclude that tobramycin improves lung function in part by reducing chronic inflammation and neutrophil-mediated lung damage in pwCF.

## Introduction

*Pseudomonas aeruginosa* is an opportunistic pathogen that infects the lungs of immunocompromised individuals, including those with chronic obstructive pulmonary disease and cystic fibrosis (CF), and is an important cause of acute pneumonia (Williams et al., 2010; Konstan et al., 2012; Lieberman and Lieberman, 2003; Novosad and Barker, 2013; Parker et al., 2016). *P. aeruginosa* is one of the leading causes of nosocomial infections worldwide, and ventilator-associated pneumonia mortality can be as high as 30% in some institutions (Williams et al., 2010). *P. aeruginosa* contributes to 5–10% of the acute exacerbations in COPD, afflicting 24 million Americans, and is the 3^rd^-leading cause of death in the U.S. (Novosad and Barker, 2013; Sethi, 2010; Lieberman and Lieberman, 2003). *P. aeruginosa* also chronically colonizes the lungs of ∼70-80% of adults with CF, and its presence is strongly associated with reduced forced expiratory volume (FEV_1_) and a progressive loss of lung function (Stanton, 2017; Davies and Martin, 2018; Cohen and Prince, 2012)

CF is a genetic disease caused by absent or aberrant function of the cystic fibrosis transmembrane conductance regulator (CFTR), which leads to airway periciliary dehydration, increased mucus viscosity, and decreased mucociliary clearance (Stanton, 2017; Stoltz et al., 2015). Insufficient mucociliary clearance causes persistent bacterial infection, non-resolving lung inflammation, and excessive neutrophil recruitment (Hauser et al., 2011; Lin and Kazmierczak, 2017). Chronic neutrophilic airway inflammation damages the lungs by continuous secretion of reactive oxygen species (ROS) and proteases, contributing to bronchiectasis and progressive CF lung function loss (Roesch et al., 2018; Khan et al., 2019). *Pseudomonas aeruginosa* is the most common pathogen identified in adult CF lungs (Hauser et al., 2011; Cystic Fibrosis Foundation, 2020), and *P. aeruginosa* respiratory infection correlates with CF lung disease severity and mortality (Emerson et al., 2002; Robinson et al., 2009).

Inhaled tobramycin is a commonly used antibiotic to suppress *P. aeruginosa* burden in CF and to ameliorate lung function loss once chronic pulmonary colonization is established (Cystic Fibrosis Foundation, 2020; Davies and Martin, 2018). The long-term use of inhaled tobramycin significantly improves lung function and reduces mortality in people with CF (pwCF) (Sawicki et al., 2012; Bowman, 2002). Inhaled tobramycin is administered in intermittent repeated cycles of 28 days on the drug and 28 days off. In a double-blind, placebo-controlled study, lung function improved significantly after the first two weeks of treatment and correlated with a decrease of *P. aeruginosa* colony-forming units (CFUs) in sputum by more than 158-fold (Ramsey et al., 1999). Intriguingly, the magnitude of the reduction in bacterial CFUs was less than ten-fold in the third cycle of therapy, although lung function improvement was maintained at a comparable level (Ramsey et al., 1999). Furthermore, an open-label, follow-on trial with adolescent patients and 12 treatment cycles revealed that the reduction of *P. aeruginosa* CFUs in sputum only explained 11.7% of CF lung function improvement (Moss, 2002). Moreover, a more recent analysis of sputum revealed that tobramycin has no significant effect on *P. aeruginosa* abundance (Nelson et al., 2020). Together, these data suggest that tobramycin improves CF lung function by an unknown mechanism in addition to its bactericidal activity. The goal of this study is to elucidate this mechanism.

In the CF lungs, *P. aeruginosa* suppresses the host immune response by secreting outer membrane vesicles (OMVs), which fuse with host cells and deliver virulence factors, DNA, small RNAs (sRNAs), and transfer RNA (tRNA) fragments that mediate inter-kingdom host-pathogen interaction (Li and Stanton, 2021; Kaparakis-Liaskos and Ferrero, 2015; Bomberger et al., 2009; Coelho and Casadevall, 2019). OMVs are 50-300 nm lipopolysaccharide (LPS)-decorated vesicles secreted by all gram-negative bacteria (Coelho and Casadevall, 2019; Jan, 2017). Recently, we reported that *P. aeruginosa* secretes a 24-nucleotides (nt) long sRNA in OMVs, which diffuse through the airway mucus layer and fuse with bronchial epithelial cells to transfer the sRNA (Koeppen et al., 2016). The sRNA down-regulates the OMV-induced secretion of IL-8, a potent neutrophil attractant, by airway epithelial cells, leading to attenuated recruitment of neutrophils into mouse lungs (Koeppen et al., 2016).

This study aimed to test the hypothesis that tobramycin prevents the decline in lung function of pwCF by increasing the level of anti-inflammatory sRNAs in OMVs secreted by *P. aeruginosa*. Here, we demonstrate that tobramycin increases the abundance of two 5′ formyl-methionine tRNA (tRNA-fMet) halves in OMVs, and that the 5′ tRNA-fMet halves are delivered into primary CF-HBECs by OMVs. 5′ tRNA-fMet halves suppress IL-8 secretion by CF-HBECs and reduce KC (a murine homolog of IL-8) levels and neutrophil recruitment in mouse lungs by an AGO2-mediated post-transcriptional regulatory mechanism. This 5′ tRNA-fMet halves-mediated reduction in lung neutrophils is predicted to mitigate lung damage. In pwCF, the IL-8 concentration and neutrophil content in bronchoalveolar lavage fluid (BALF) was significantly reduced during the month of tobramycin administration compared to the month off tobramycin. Taken together, these data reveal that the clinical benefit of tobramycin is due in part to an increase in the secretion of 5′ tRNA-fMet halves in OMVs, leading to attenuation of IL-8 and neutrophil-mediated CF lung damage.

## Results

### Tobramycin reduces the ability of OMVs secreted by *P. aeruginosa* to stimulate IL-8 secretion by CF-HBECs

To test the hypothesis that tobramycin alters the virulence of OMVs secreted by *P. aeruginosa*, we designed an *in vitro* experiment depicted in Figure 1A. *P. aeruginosa* strain PA14 was grown in lysogeny broth (LB), and OMVs secreted by *P. aeruginosa* treated with vehicle (ctrl-OMVs) or tobramycin (Tobi-OMVs) were isolated as described in Methods. The concentration of tobramycin used (1 μg/mL) reduced growth by 33%, an amount similar to that observed in pwCF treated with tobramycin after three cycles of therapy (Ramsey et al., 1999) (Figure 1B). Tobramycin increased the secretion of OMVs by 38% compared to control (Figure 1C), which coincides with a previous report that antibiotics induce OMV production by *P. aeruginosa* (MacDonald and Kuehn, 2013).

**Figure 1.**
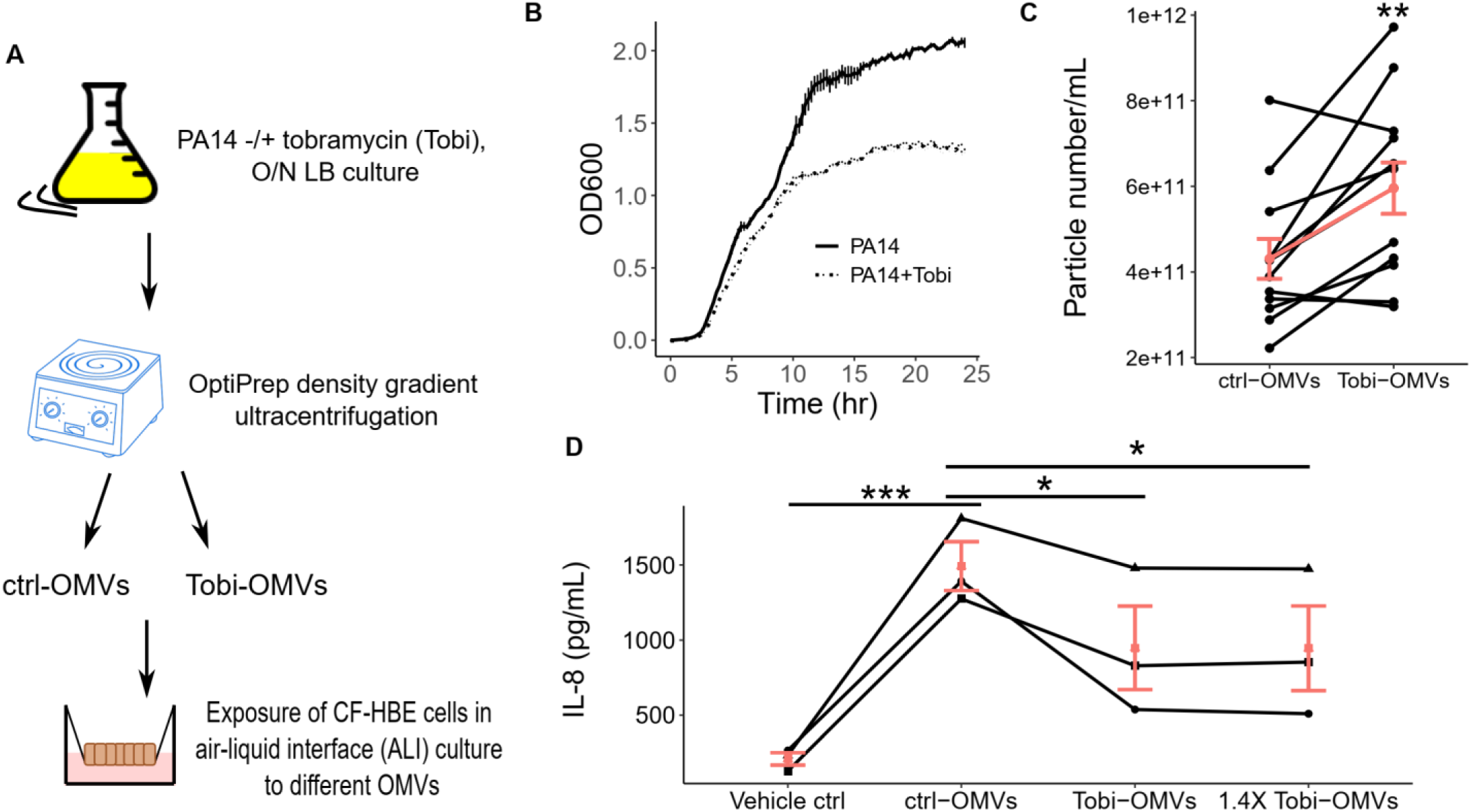
Tobramycin reduces the ability of OMVs secreted by *P. aeruginosa* to stimulate IL-8 secretion by CF-HBECs. (**A**) A schematic diagram shows the experimental design. (**B**) Growth curve (in microtiter plates) of PA14 in LB alone (PA14) or in LB with tobramycin (1 μg/mL; PA14+Tobi). Lines represent the averages from three biological replicates, and error bars indicate standard error of means (SEM). The presence of tobramycin inhibited PA14 growth by 33% between the 13 to 20 hour time points. (**C**) OMV concentration of purified ctrl-OMVs and Tobi-OMVs (*n* = 12) measured with nanoparticle tracking analysis (Nanosight NS300). The red line connects the mean concentration of the two groups and demonstrates a 38% increase in Tobi-OMVs concentration compared to ctrl-OMVs concentration. Data are shown as the means ± SEM (**D**) Primary CF-HBECs from three donors (*n* = 3) were polarized in ALI culture before being exposed to either the same number of ctrl-OMVs or Tobi-OMVs or 40% more Tobi-OMVs (1.4X Tobi-OMVs) for 6 hours. The basolateral medium was collected to measure IL-8. Lines connect experiments conducted with CF-HBECs from the same donor. Horizontal red lines and red dots indicate means ± SEM. Paired t-tests (**C**); Linear mixed-effects models with CF-HBEC donor as a random effect were used to calculate *P* values (**D**); \**P* < 0.05; \*\**P* < 0.01;\*\*\**P* < 0.001.

To examine the effect of OMVs on the host immune response, polarized HBECs from CF donors (CF-HBECs) were grown in air-liquid interface (ALI) culture (Randell et al., 2011; Barnaby et al., 2019) and exposed to the same number of ctrl-OMVs or Tobi-OMVs for 6 hours, whereupon IL-8 secretion was measured. Tobi-OMVs induced 36% less IL-8 secretion than ctrl-OMVs (Figure 1D). Similar results were obtained even when cells were exposed to 40% more Tobi-OMVs (1.4X Tobi-OMVs) than ctrl-OMVs to reflect the finding that tobramycin increased OMV production (Figure 1D). We measured other cytokines secreted by CF-HBECs (EGF, GRO, IL17a, and IP10) in response to ctrl-OMVs and Tobi-OMVs, but of these additional cytokines only IP-10 was significantly reduced by Tobi-OMVs compared to ctrl-OMVs (Supplemental Figure 1).

### Tobramycin increases the abundance of 5′ tRNA-fMet halves in OMVs, and the tRNA halves are transferred from OMVs to CF-HBECs

To test the hypothesis that tobramycin increases the abundance of anti-inflammatory sRNAs in OMVs, we performed a small RNA-sequencing analysis to compare the sRNA content in ctrl-OMVs and Tobi-OMVs. We identified 6145 unique sequences mapped to the PA14 genome, and 1064 were differentially enriched in Tobi-OMVs. The sequence length ranged from 20 to 48 nucleotides; however, we excluded the 48-nt sequences from further analysis as they represented RNA species longer than the read length.

We focused on the most abundant and differentially induced sRNAs in Tobi-OMVs (Figure 2A and Table 1). We chose two 35-nt long sRNAs (#5 and #7 in Table 1) that were fragments of two initiator tRNAs (tRNA-fMet1 and tRNA-fMet2 located at PA14_62790 and PA14_52320, respectively) in PA14 for further analysis because they were bioinformatically predicted to suppress IL-8 secretion by CF-HBECs. The sequence reads in Tobi-OMVs mapped to these two loci had similar length distributions (Figure 2B and 2C; 80% of reads were 35-nt long), suggesting the tRNA-fMet fragments were not products of random degradation. The similar length distributions also imply common machinery for the biogenesis of these two tRNA fragments. Both have low minimum free energy, suggesting stable secondary structures.

**Table 1.** Top 10 most abundant and most differentially induced sRNAs in Tobi-OMVs compared to ctrl-OMVs.

| # | PA14 locus | Gene product of the locus | Log2FC | Average Log2CPM | length | Minimum free energy (kcal/mol)<sup>A</sup> |
| --- | --- | --- | --- | --- | --- | --- |
| 1 | Multiple | tRNA-Asp | 4.95 | 10.93 | 23 (20, 23)<sup>B</sup> | -0.2 |
| 2 | Multiple | 16S rRNA | 3.98 | 10.83 | 33 (30-39)<sup>B</sup> | -6.9 |
| 3 | Multiple | 23S rRNA | 5.57 | 10.35 | 44 | -14.6 |
| 4 | Multiple | tRNA-Ala | 3.19 | 10.13 | 34 | -8 |
| 5 | 62790 | tRNA-fMet1 | 4.82 | 9.88 | 35 | -7.5 |
| 6 | 28740 | tRNA-Pro | 2.70 | 9.39 | 36 | -9.7 |
| 7 | 52320 | tRNA-fMet2 | 2.74 | 8.95 | 35 | -9.3 |
| 8 | 61760 | tRNA-Gln | 6.23 | 8.88 | 20 | -0.2 |
| 9 | 30720 | tRNA-Cys | 3.41 | 8.85 | 40 | -4.6 |
| 10 | multiple | 5S rRNA | 2.72 | 8.75 | 45 | -3.8 |

<sup>A</sup>The minimum free energy for each sRNA was predicted using the RNAfold web server (Gruber et al., 2008)
<sup>B</sup>Multiple reads of different lengths were mapped to the same locus, and the most abundant read is listed in Table 1 and Figure 2A.

**Figure. 2.**
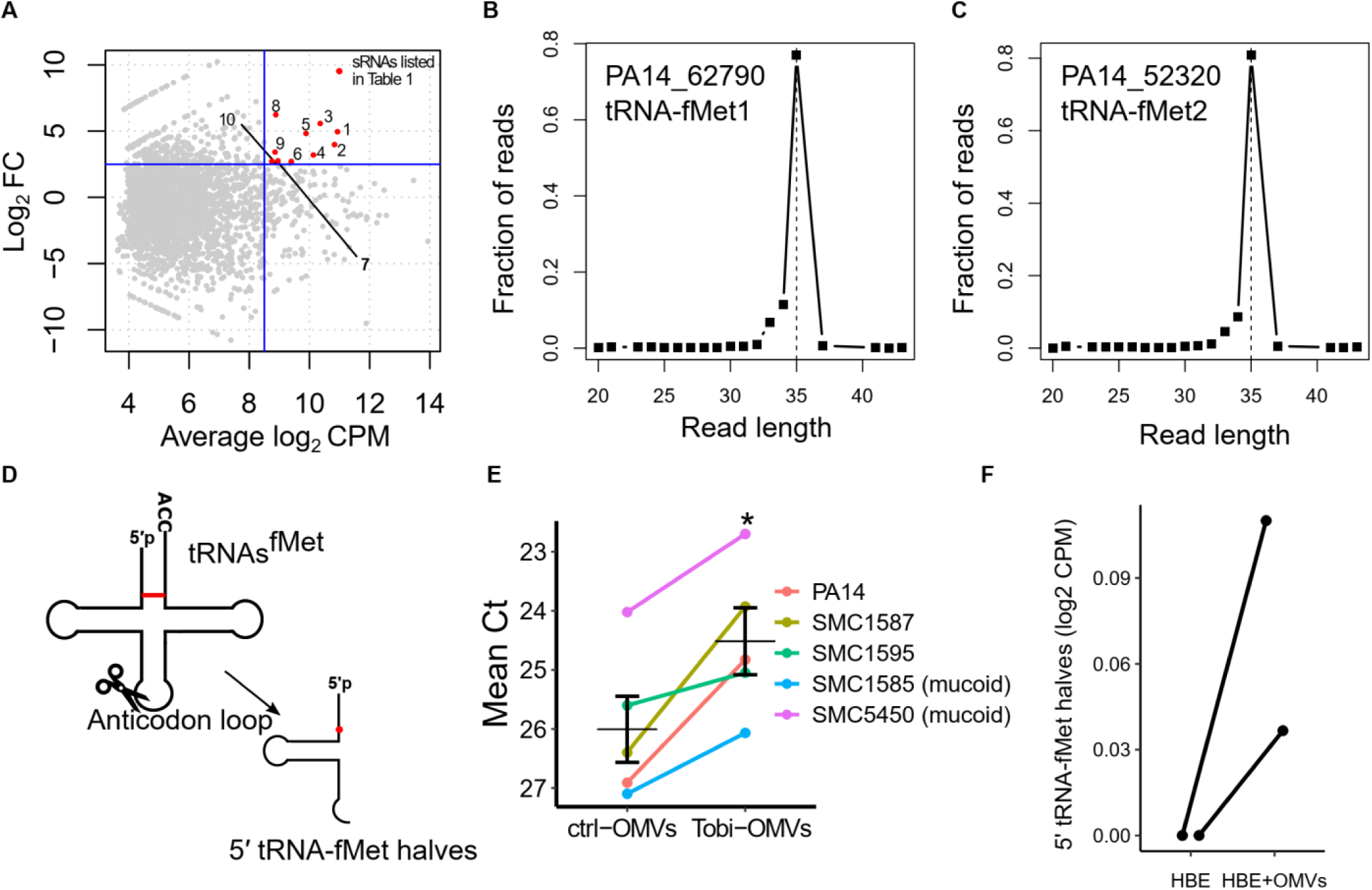
Tobramycin increases the abundance of 5′ tRNA-fMet halves in OMVs, and the tRNA halves are transferred into host cells. (**A**) MA plot comparing the small RNA expression profile in Tobi-OMVs and ctrl-OMVs (*n* = 3 for each group). Each dot represents a unique sequence read. The most abundant and most induced sRNAs by tobramycin treatment are highlighted in red and listed in Table 1. (**B and C**) Length distribution of Tobi-OMVs sRNAs mapped to gene locus PA14_62790 (**B**) and PA14_52320 (**C**). (**D**) Secondary cloverleaf structure of tRNAs^fMet^ and cleavage site in the anticodon loop to generate 5′ tRNA-fMet halves. The red line indicates the only different pair of nucleotides between the two tRNAs^fMet^, and the red dot represents the only nucleotide difference between the two 5′ tRNA-fMet halves. (**E**) qPCR for 5′ tRNA-fMet halves in ctrl-OMVs and Tobi-OMVs purified from PA14 and four clinical isolates (*n* = 5 strains), including two mucoid and two non-mucoid strains. The qPCR primers and probe were designed to detect both 5′ tRNA-fMet halves. Horizontal lines indicate means ± SEM. A paired t-test was used to establish significance. * *P* < 0.05. (**F**) Both 5′ tRNA-fMet halves were detected in polarized primary HBE cells exposed to ctrl-OMVs but not in unexposed cells using small RNA sequencing (from two donors; *n* = 2). Sequence reads in (F) are from our previously published dataset (Koeppen et al., 2016).

The two 35-nt long tRNA-fMet fragments are 5′ halves of tRNAs^fMet^ (hereafter called 5′ tRNA-fMet halves), which are products of cleavage in the anticodon loop (Figure 2D). Importantly, the two 5′ tRNA-fMet halves have high sequence similarity with only one nucleotide difference, suggesting similar sequence-based targeting functions. The high sequence similarity allowed us to design qPCR primers to quantify both 5′ tRNA-fMet halves simultaneously. By qPCR, we found that Tobi-OMVs secreted by PA14 and four clinical isolates (including two mucoid strains) also contained significantly more 5′ tRNA-fMet halves than ctrl-OMVs (Figure 2E), indicating a strain-independent phenotype that extends to clinically relevant strains. Moreover, we reanalyzed our previously published small RNA-sequencing experiment (Koeppen et al., 2016), in which we sought to detect PA14 sRNAs transferred into non-CF HBECs after ctrl-OMVs exposure, and we were able to identify both 5′ tRNA-fMet halves in OMV-exposed cells but not in the un-exposed group (Figure 2F).

Taken together, these observations demonstrate that the two 5′ tRNA-fMet halves are the most abundant and most differentially induced sRNAs in Tobi-OMVs, have high sequence similarity, and are delivered to airway epithelial cells by OMVs; thus, they are good candidates for further investigation into their possible role in suppressing IL-8 secretion.

### 5′ tRNA-fMet halves reduce IL-8 secretion

To determine if 5′ tRNA-fMet halves reduce IL-8 secretion, we transformed PA14 with an arabinose-inducible vector expressing 5′ tRNA-fMet1 half (tRNA1-OMVs) or an empty vector control (V-OMVs). Small RNA-sequencing confirmed that the expression of 5′ tRNA-fMet1 half in tRNA1-OMVs was significantly induced by 2.73 fold compared to V-OMVs (Supplemental Figure 2). Primary CF-HBECs were exposed to V-OMVs or tRNA1-OMVs, and the secretion of IL-8 was measured by ELISA. As predicted, tRNA1-OMVs induced less IL-8 secretion compared to the same amount of V-OMVs (Figure 3A). To provide additional support for the conclusion that 5′ tRNA-fMet halves reduce IL-8 secretion by CF-HBECs, we designed an inhibitor, an RNA oligonucleotide with a complementary sequence to both 5′ tRNA-fMet halves. CF-HBECs were transfected with the inhibitor or negative control inhibitor followed by exposure to ctrl-OMVs or 1.4X Tobi-OMVs. As predicted, the inhibitor reduced the ability of 1.4X Tobi-OMVs to suppress IL-8 secretion compared to ctrl-OMVs (Figure 3B).

**Figure 3.**
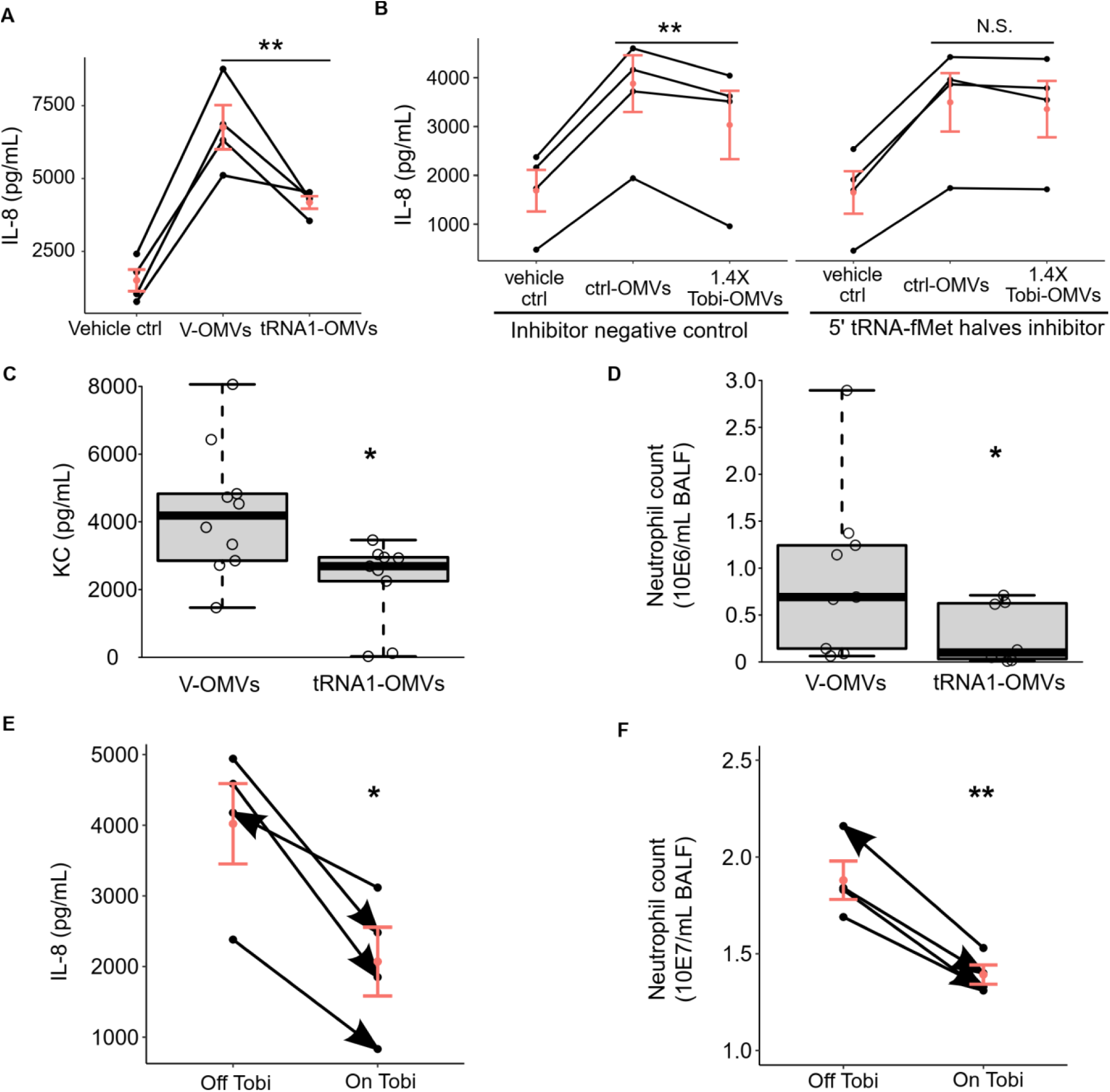
5′ tRNA-fMet halves reduce IL-8 secretion *in vitro* and *in vivo*. **(A)** Polarized CF-HBECs (*n* = 4) exposed to tRNA1-OMV secreted less IL-8 compared to cells exposed to V-OMV. (**B**) The Tobi-OMV effect of reducing IL-8 secretion was abolished by transfection of an antisense RNA oligo inhibitor that anneals to both 5′ tRNA-fMet halves (5′ tRNA-fMet halves inhibitor) but not by transfection of a negative control inhibitor (*n*= 4). Lines in panels A and B connect data points using the cells from the same donor, a biological replicate. A linear mixed-effects model with CF-HBEC donor as a random effect was used to calculate *P* values. (**C** and **D**) BALF from mice exposed to V-OMVs or tRNA1-OMVs was collected to measure KC concentration (**C**) and neutrophil number (**D**). Minimum-to-maximum whisker and box plots showing the median and interquartile ranges. 9 to 10 mice were used per group, and Wilcoxon rank-sum tests were used to test significance. (**E** and **F**) BALF samples collected from four CF subjects (*n*= 4) during the 4-week administration of inhaled tobramycin (On Tobi) or not (Off Tobi) with BALF IL-8 levels in panel (**E**) and BALF neutrophil content in panel (**F**). Lines connect data points from the same subject, and the arrowheads indicate the sample collection order for each CF subject. Linear mixed-effect models were used to account for donor-to-donor variability and the number of days between collection dates for each sample pair (Supplemental Table 1). Horizontal red lines and red dots indicate means ± SEM; N.S., not significant; \**P* < 0.05; \*\**P* < 0.001.

Studies were conducted in mice to further support the conclusion that tobramycin reduces the pro-inflammatory effect of OMVs by increasing the 5′ tRNA-fMet1 half content. Mice were exposed to the same number of V-OMVs or tRNA1-OMVs by oropharyngeal aspiration for 5 hours, and BALF samples were harvested for analysis. The concentration of KC, a murine functional homolog of IL-8 (Figure 3C), and neutrophil content (Figure 3D) were significantly reduced in BALF obtained from mice exposed to tRNA1-OMVs compared to V-OMVs. Thus, 5′ tRNA-fMet1 half reduced the pro-inflammatory response of CF-HBECs *in vitro* and in a mouse model of inflammation.

### Inhaled tobramycin has an anti-inflammatory effect in *P. aeruginosa*-infected CF lungs

To determine if tobramycin reduces inflammation and neutrophil burden in CF lungs, we performed a retrospective analysis to assess whether the administration of inhaled tobramycin changes the inflammatory status in pwCF. BALF samples were collected from four CF patients chronically infected with *P. aeruginosa* during the month of inhaled tobramycin (On Tobi) and the month off tobramycin (Off Tobi). In BALF obtained On Tobi, average IL-8 levels were reduced by 48.5% (Figure 3E), and the number of neutrophils was decreased by 25.9% (Figure 3F) compared to Off Tobi. This clinical observation is consistent with the *in vitro* and mouse experiments, suggesting that OMVs secreted by tobramycin-exposed *P. aeruginosa* are less pro-inflammatory than control OMVs.

### 5′ tRNA-fMet halves regulate gene expression by base-pairing with target genes in CF-HBECs using an AGO2-mediated mechanism

Although we and others have shown that prokaryotic sRNAs regulate eukaryotic gene expression in a sequence-specific manner (Koeppen et al., 2016; Maute et al., 2013), the mechanism is unknown. We therefore conducted experiments to determine if *P. aeruginosa* 5′ tRNA-fMet halves can utilize the eukaryotic Argonaute 2 (AGO2) dependent gene silencing complex to suppress IL-8 secretion. We designed a three-step approach to identify the RNA binding targets, followed by proteomic analysis to determine the effect of 5′ tRNA-fMet halves on protein expression (Figure 4A). Ingenuity Pathway Analysis (IPA) (Krämer et al., 2014) was performed at each step to identify significantly enriched and down-regulated pathways that are relevant to CF and predicted to decrease IL-8 secretion (Table 2).

**Figure 4.**
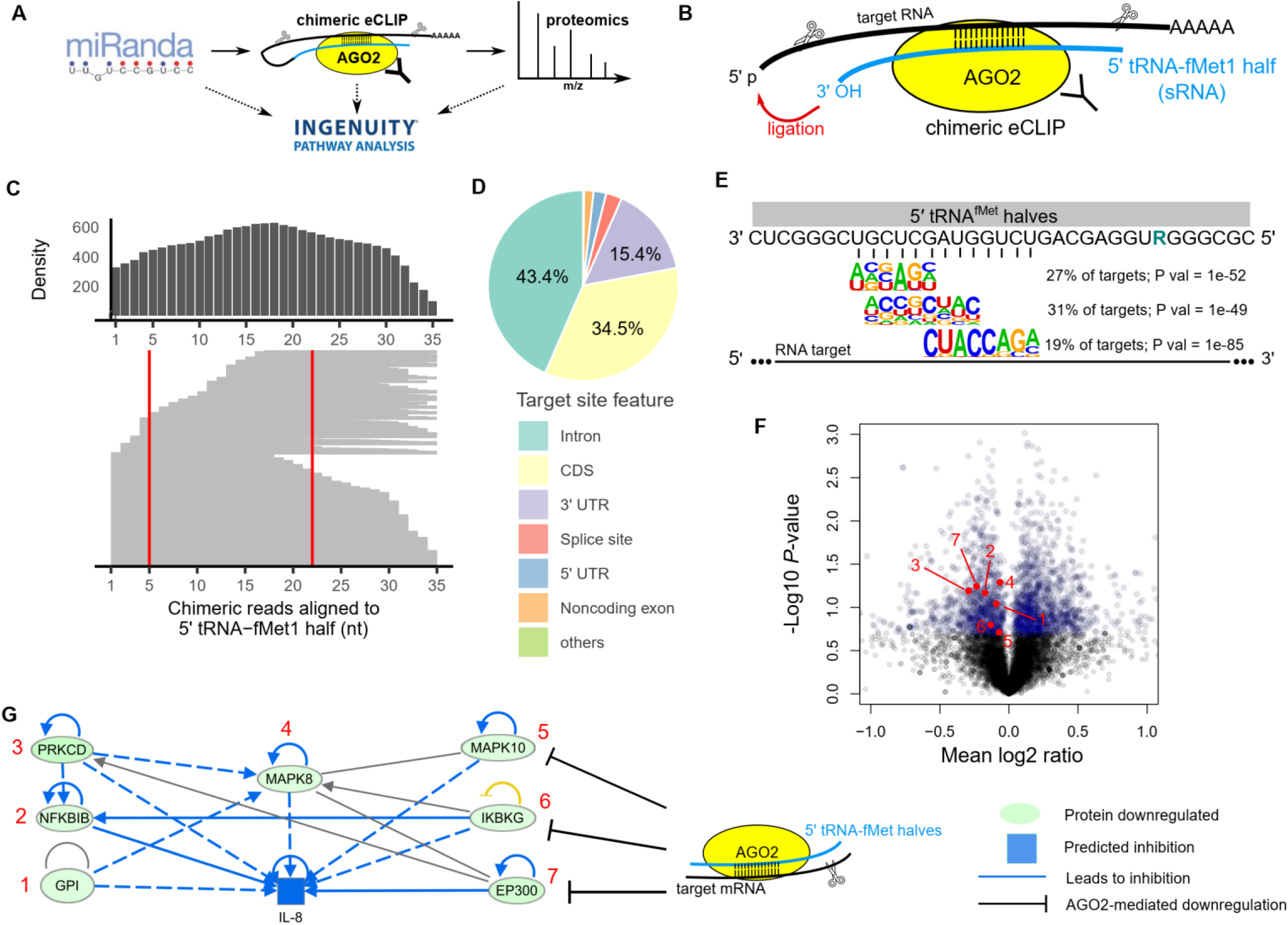
5′ tRNA-fMet halves downregulate protein expression in a sequence-specific manner mediated by AGO2. (**A**) Schematic representation of the three-step approach to identify targets of 5′ tRNA-fMet halves leading to differential protein expression. (**B**) Diagram of chimeric eCLIP to identify RNAs pulled down with AGO2. Samples were treated with RNase I for RNA fragmentation followed by immunoprecipitation of AGO2-mRNA-sRNA complexes before mRNA-sRNA ligation to generate chimeric reads. (**C**) Alignment of chimeric reads to the 5′ tRNA-fMet1 half sequence and the count distribution of each nucleotide (top density plot). Chimeric reads were identified in tRNA-fMet1 half-transfected CF-HBECs (one donor; *n* = 1) and contained at least 18-nt long 5′ tRNA-fMet1 half subsequences. The red lines indicate the region where the primer performs targeted chimeric eCLIP anneals. (**D**) Distribution of target site features identified with the targeted chimeric eCLIP. (**E**) Sequence logos of most significant enriched target RNA motifs and the complementary sequence of 5′ tRNA-fMet halves. R denotes a purine nucleotide (G/A). (**F**) Volcano plot of proteomic analysis of polarized CF-HBECs (three donors; *n* = 3) treated with tRNA1-OMVs compared to cells treated with V-OMVs. The top 20% differentially expressed proteins, determined by paired t-tests, are colored in blue. Red dots with numbers represent down-regulated proteins corresponding to proteins numbered in panel (G). (**G**) IPA identified a down-regulated pro-inflammatory network in the five consensus pathways (table 2), leading to decreased IL-8 expression. mRNA transcripts encoding MAPK10, IKBKG, and EP300 were identified as binding targets of tRNA-fMet1 half in the targeted chimeric eCLIP experiment.

**Table 2.** The consensus of significantly enriched signaling pathways identified using three approaches^A^.

| Canonical pathway | Gene target prediction |  | Gene target validation |  | Protein expression |  |
| --- | --- | --- | --- | --- | --- | --- |
|  | miRanda (1518)<sup>B</sup> |  | chimeric eCLIP (1936)<sup>B</sup> |  | Proteomics (2168)<sup>B</sup> |  |
|  | *P* value | z-score<sup>C</sup> | *P* value | z-score<sup>C</sup> | *P* value | z-score<sup>C</sup> |
| **Integrin-linked kinase (ILK) Signaling** | 0.0005 | -3.26 | 0.0041 | -0.53 | 1.1E-08 | -2.27 |
| **LPS-stimulated MAPK Signaling** | 0.0087 | -3.46 | 0.0005 | -3.60 | 7.7E-05 | -0.68 |
| **HIF1a Signaling** | 0.0022 | -3 | 0.0072 | -4.35 | 0.0015 | -0.87 |
| **IL-17A Signaling in Airway Cells** | 0.0011 | -2.53 | 0.0085 | -1.66 | 0.0077 | -0.30 |
| **IL-6 Signaling** | 0.0002 | -4.02 | 0.0389 | -3.46 | 0.0141 | -0.94 |

<sup>A</sup> There are 38 consensus pathways. Only consensus pathways predicted to downregulate IL-8 secretion in epithelial cells by IPA are listed.
<sup>B</sup> Number in parentheses indicates the number of genes/proteins used to perform pathway enrichment analysis.
<sup>C</sup> Negative z-scores indicate that the pathways are predicted to be down-regulated.

We first performed a miRanda microRNA (miRNA) target scan (Enright et al., 2003) to predict the human binding targets of 5′ tRNA-fMet1 half. miRanda is an algorithm designed for RNA-RNA binding prediction considering sequence complementarity and binding free energy. Given the high sequence similarity between the two sRNAs, we used the sequence of 5′ tRNA-fMet1 half to scan the whole human transcriptome and adjusted the prediction for the gene expression profile of polarized HBE cells to identify a list of 1518 predicted targets, accounting for 8.4% of human coding genes. IPA identified several pro-inflammatory pathways in epithelial cells that are predicted to be down-regulated by 5′ tRNA-fMet1 half, including integrin-linked kinase signaling (Eucker et al., 2014; Ahmed et al., 2014; Gravelle et al., 2010), LPS-stimulated MAPK Signaling (Koeppen et al., 2016), and HIF1a signaling (Cane et al., 2010) (Table 2).

The target gene prediction and pathway analysis encouraged us to map transcriptome-wide interactions between 5′ tRNA-fMet1 half and target mRNAs mediated by AGO2. Although tRNA fragments have been shown to regulate gene expression, to our knowledge, there is no direct evidence that tRNA halves or any small noncoding RNA secreted by a prokaryotic organism can suppress eukaryotic gene expression by interacting with the AGO2 gene silencing complex. Thus, to determine if 5′ tRNA-fMet1 half interacts with eukaryotic mRNAs in the AGO2-containing complex, we utilized the enhanced crosslinking and immunoprecipitation (eCLIP) approach (Van Nostrand et al., 2016). Briefly, and as described in detail in methods, this approach involved transfection of 5′ tRNA-fMet1 half into CF-HBECs, followed by ligation of 5′ tRNA-fMet1 half and other small RNAs to target mRNAs yielding sRNA-mRNA chimeric fragments. Chimeric fragments immunoprecipitated with AGO2 were characterized with high-throughput sequencing (chimeric eCLIP) (Figure 4B), which provided an unprecedented resolution to identify sRNA-mRNA interactions. CF-HBECs were transfected with 5′ tRNA-fMet1 half or negative control siRNA (siNC) and subjected to AGO2 chimeric eCLIP analysis. This analysis allowed us to profile transcriptome-wide AGO2 authentic binding sites mediated by 5′ tRNA-fMet1 half and other miRNAs with a stringent cutoff (IP vs. input cluster log2 fold enrichment ≥ 3 and *P* value ≤ 0.001). Using these parameters, we identified 10947 AGO2 binding sites in 4454 genes. Within those authentic binding sites, we identified 629 chimeric reads containing at least 18-nt long subsequences of 5′ tRNA-fMet1 half. Lengths of identified subsequences ranged from 18-nt to the full size of 5′ tRNA-fMet1 half, and alignment positions of subsequences were evenly distributed (Figure 4C). These findings suggest that the full length 5′ tRNA-fMet1 half was loaded into the AGO2 complex for gene targeting without being pre-processed into a shorter sRNA, and the subsequences with different lengths identified in chimeric reads were products of RNA fragmentation, a key step in the eCLIP sequencing library preparation.

To deeply profile the target repertoire of 5′ tRNA-fMet1 half mediated by AGO2, we designed a 5′ tRNA-fMet1 half specific primer, which anneals to most of the identified subsequence (Figure 4C), for targeted sequencing (targeted chimeric eCLIP). The targeted chimeric eCLIP allowed us to sequence 5′ tRNA-fMet1 half containing chimeric fragment at a much higher depth. We identified that 5′ tRNA-fMet1 half targeted 5776 sites in 1945 genes, and those target sites were not found in the negative control (siNC) transfected cells. Interestingly, although miRNA-AGO2 complexes usually target 3′ untranslated regions (3′ UTR), most 5′ tRNA-fMet1 half-AGO2 target sites were in introns (Figure 4D). Furthermore, motif enrichment analysis in target sites revealed that nucleotides 16-28 from the 5’-end of 5′ tRNA-fMet1 half (which do not include the only distinct nucleotide between the two 5′ tRNA-fMet halves) could explain 77% of identified target sites (Figure 4E). The fact that the most popular binding motif does not contain the unique nucleotide differentiating the two 5′ tRNA-fMet halves suggests that they have many common target genes. Moreover, we found that miRanda predicted the target genes identified by chimeric eCLIP significantly better than expected by chance (Fisher′ s exact test, *P* < 10^−12^), suggesting that bioinformatic target prediction methods based on base-pairing can reliably predict target genes. Also, IPA predicted that a similar set of pro-inflammatory and IL-8 induction pathways were inhibited by down-regulating these target genes (Table 2).

Lastly, to identify proteins whose abundances were changed by 5′ tRNA-fMet halves delivered by OMVs, we utilized OMVs secreted by the 5′ tRNA-fMet1 half-overexpression and empty vector clones (Supplemental Figure 2). Primary CF-HBECs from three donors were exposed to V-OMVs or tRNA1-OMVs for 6 hours before being subjected to proteomic analysis. 8343 proteins were identified, and we selected the top 20% differentially expressed proteins by *P* value, yielding 943 down-regulated proteins (Figure 4F). The statistically enriched and down-regulated pathways identified by IPA overlapped with our previous analysis based on identified target genes (Table 2). These down-regulated pathways included downstream signaling of IL17A and IL-6, which are pro-inflammatory cytokines secreted by other cell types in CF lungs to induce IL-8 secretion by CF-HBECs (McAllister et al., 2005; Courtney et al., 2004; Hsu et al.). Considering the top 20% down-regulated proteins, IPA identified seven proteins that contributed to IL-8 expression and predicted the decrease of IL-8 secretion (Figure 4G). Among the seven significantly down-regulated proteins, MAPK10, IKBKG, and EP300 were the direct targets of 5′ tRNA-fMet1 half identified with our targeted chimeric eCLIP experiment, suggesting targeting of a pro-inflammatory network involving MAPK and NFκB signaling.

In summary, our three-step approach demonstrated that 5′ tRNA-fMet1 halves transferred from OMVs to CF-HBECs were loaded into the AGO2 complex to target specific genes via a base-pairing mechanism, thus mediating the Tobi-OMVs induced reduction in IL-8 secretion.

## Discussion

The goal of this study was to determine how tobramycin improves clinical outcomes in pwCF without significantly reducing the abundance of *P. aeruginosa*. Our data reveal that tobramycin increases the concentration of 5′ tRNA-fMet halves in OMVs secreted by *P. aeruginosa*, that the OMVs deliver 5′ tRNA-fMet halves to CF-HBECs, and that the increased delivery of 5′ tRNA-fMet halves to CF-HBECs suppresses IL-8 secretion by interacting with pro-inflammatory gene transcripts, including MAPK10, IKBKG, and EP300, in an AGO2-mediated mechanism. Both *in vitro* and *in vivo* experiments in mice are consistent with this conclusion. Moreover, our retrospective analysis of pwCF on and off tobramycin is consistent with our data in mice that tobramycin reduces IL-8 and the neutrophil content in BALF. The reduction in the neutrophil content in BALF is predicted to mitigate lung damage in the CF lungs since CF neutrophils are the source of significant lung damage in pwCF (Figure 5). To our knowledge, this is the first report demonstrating that tRNA halves secreted by a prokaryotic organism suppress gene expression in eukaryotic cells by an AGO2-mediated RNA silencing mechanism.

**Figure 5.**
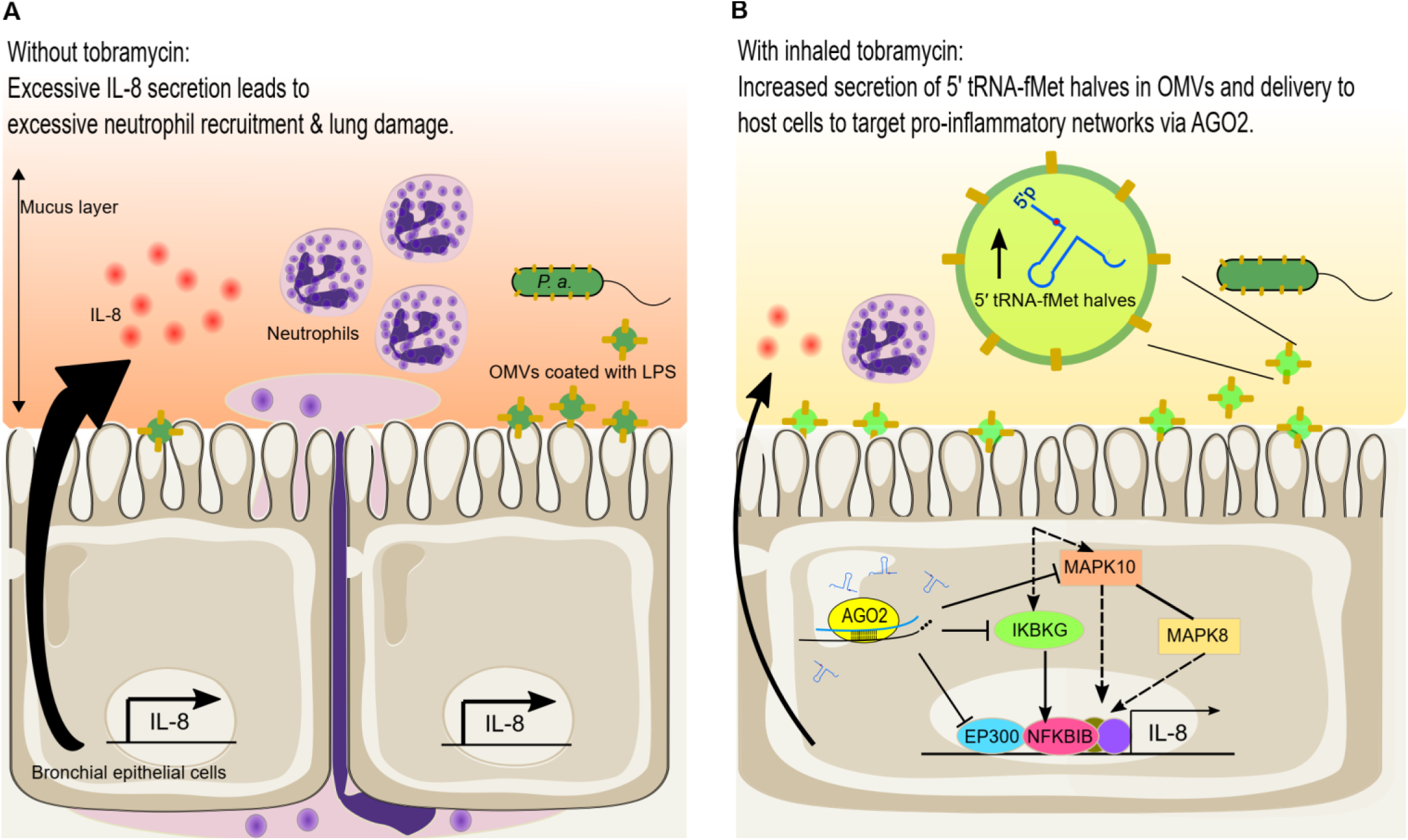
Graphical abstract indicating the anti-inflammatory effect of tobramycin mediated by 5′ tRNA-fMet halves in *P. aeruginosa* OMVs. **(A)** *P. aeruginosa* colonizes the CF lungs and secrets OMVs. OMVs diffuse through the mucus layer overlying bronchial epithelial cells and induce IL-8 secretion, which recruits excessive neutrophils and causes lung damage. (**B**) Tobramycin increases 5′ tRNA-fMet halves in OMVs secreted by *P. aeruginosa*. 5′ tRNA-fMet halves are delivered into host cells and loaded into the AGO2 protein complex to down-regulate protein expression of MAPK10, IKBKG, and EP300, which suppresses OMV-induced IL-8 secretion and neutrophil recruitment. A reduction in neutrophils in BALF is predicted to improve lung function and decrease lung damage.

tRNA-derived fragments are a novel class of regulatory sRNAs in prokaryotes and eukaryotes (Li and Stanton, 2021; Su et al., 2020). tRNAs are the most abundant RNA species by the number of molecules, and fragments with different lengths have been reported in three domains of life. miRNA-sized (∼24-nt long) tRNA fragments in mammalian cells have garnered attention as they have been found to associate with Argonaute (AGO) proteins to mediate gene silencing by base-pairing with target mRNAs (Maute et al., 2013; Kumar et al., 2014; Kuscu et al., 2018). Recently, a report showed that in *Bradyrhizobium japonicum*, a 21-nt tRNA fragment utilizes host plant AGO1 to regulate host gene expression, a cross-kingdom symbiotic relationship between bacteria and plants (Ren et al., 2019). However, tRNA halves have been thought not to involve the AGO-miRNA-like mechanism since they are 32–50 nt in length. tRNA halves have been shown to have both positive and negative effects on translation by regulating the formation of ribosomes and the translation initiation complex (Ivanov et al., 2011; Gebetsberger et al., 2012; Fricker et al., 2019; Mleczko et al., 2018). Here, we provide the first evidence that 35-nt 5′ tRNA-fMet halves from a bacterial pathogen are transferred into eukaryotic cells and loaded into AGO2-containing protein complexes without pre-processing into shorter sRNAs to suppress gene expression by base-pairing with target mRNAs. This finding not only provides a novel role of tRNA halves in host-pathogen interactions but also suggests that siRNAs may be an effective therapeutic approach to reduce lung damage caused by chronic inflammation and excessive neutrophil recruitment.

The production of tRNA halves, also called tRNA-derived stress-induced RNAs (tiRNAs), by cleavage in the anticodon loop of mature tRNAs is conserved across all life domains in response to various stress conditions (Tao et al., 2020; Fricker et al., 2019; Thompson et al., 2008); however, the molecular mechanism by which prokaryotes sense the stress and cleave specific tRNAs, and the cell-autonomous effects of tRNA halves induced by stress remain largely unknown. *Mycobacterium tuberculosis* maintains its persistence in host cells by cleaving several tRNAs in half with endonucleases VapCs and MazF-mt9 to reduce the level of translation (Winther et al., 2016; Schifano et al., 2016). *Escherichia coli* secretes colicin D, an anticodon ribonuclease (ACNase), to cleave tRNA-Arg of competing *E. coli* strains in half by recognizing the anticodon-loop sequence, and the cleaved tRNA-Arg blocks the ribosome A-site to disrupt translation (Tomita et al., 2000; Ogawa et al., 2020). Similar mechanisms have been identified in fungi to inhibit the cell growth of nonself competitors (Chakravarty et al., 2014; Lu et al., 2005). Together, these reports and our findings suggest that *P. aeruginosa*, in response to tobramycin exposure, up-regulates an unidentified ACNase that recognizes and cleaves the anticodon loop of tRNA-fMets, which are essential initiator tRNAs, to slow down cell growth and prevent cell death.

The interaction of *P. aeruginosa* 5′ tRNA-fMet halves with CF-HBEC mRNAs by base-pairing is reminiscent of miRNA-mRNA interactions in mammalian cells. While canonical miRNAs use a seed region, typically nucleotides 2-7 of miRNAs, to base-pair with the 3′ UTR of target mRNAs (Chipman and Pasquinelli, 2019), we observed that tRNA-fMet halves used nucleotides 16-28 to bind introns and coding sequences of target genes. This finding suggests that more studies are needed to better understand the targeting rule of tRNA halves for more accurate target prediction. Given the high sequence similarity of tRNAs in prokaryotes (Saks and Conery, 2007), precise target predictions would help generalize experimental findings to other fragments or pathogens. For example, *Helicobacter pylori* secretes sR-2509025, a 31-nt 5′ tRNA-fMet fragment, in OMVs that fuse with human gastric adenocarcinoma cells and sR-2509025 diminishes LPS-induced IL-8 secretion (Zhang et al., 2020). Due to the high sequence similarity of tRNA-fMets from *P. aeruginosa* and *H. pylori* and the similar phenotype on regulating IL-8 secretion, sR-2509025 may interact with AGO2 in gastric epithelial cells to target the pro-inflammatory network identified in this study.

Numerous reports have demonstrated that a myriad of signaling pathways, including NF-κB and MAPK signaling pathways, induce IL-8 secretion (Lee et al., 2018; Li et al., 2002, 8; Jundi and Greene, 2015). IKBKG, also known as NF-κB essential modulator (NEMO), is critical for NF-κB pathway activation. Furthermore, EP300, also called P300, is a transcription co-factor required for NF-κB-dependent IL-8 induction (Berghe et al., 1999; Huang et al., 2017). Moreover, a study demonstrated that DNA damage leads to NF-κB activation followed by MAPK10-mediated IL-8 secretion (Biton and Ashkenazi, 2011). Indeed, the elevated DNA damage response correlates with the non-resolving neutrophilic inflammation in the CF airways (Brown et al., 1995; Fischer et al., 2013); hence, our findings revealing that 5′ tRNA-fMet halves-AGO2 complex decrease IKBKG, EP300, and MAPK10 protein expression and thereby reduce IL-8 secretion and neutrophil levels are consistent with the literature.

CFTR is a negative regulator of the pro-inflammatory response mediated by MAPK and NF-κB signaling. Studies have shown that impaired CFTR leads to overactivation of NF-κB signaling and enhanced secretion of IL-8 by epithelial cells (DiMango et al., 1998; Vij et al., 2009). Also, CFTR down-regulates thermal injury-induced MAPK/NF-κB signaling, a pathway that leads to IL-8 expression and pulmonary inflammation (Dong et al., 2015). Here, we demonstrate that 5′ tRNA-fMet halves target a pro-inflammatory network involving the MAPK and NF-κB signaling pathways, which are intrinsically over-activated in CF, highlighting the importance of this network in pulmonary inflammation.

There are a few limitations of our study. First, we performed a retrospective analysis of BALF samples collected from pwCF on and off inhaled tobramycin; however, we could not collect BALF in consecutive months on and off tobramycin in the same individuals because of the invasive nature of the technique and IRB restrictions on research bronchoscopies at Dartmouth-Hitchcock Medical Center. Nevertheless, after adjusting the number of days between collection dates for each sample pair (range from 175 to 791 days; Supplemental Table 1), tobramycin-on BALF had significantly lower IL-8 concentration and fewer neutrophil counts than tobramycin-off BALF. Importantly, a similar observation was made for IL-8 in CF sputum samples collected in consecutive months from pwCF on and off tobramycin (Husson et al., 2005). Because studies have shown that IL-8 concentration in sputum is inversely correlated with pulmonary function (Sagel et al., 2002, 2001), we conclude that inhaled tobramycin has an anti-inflammatory effect in *P. aeruginosa*-infected CF lungs, resulting in improved lung function. Second, since there are many other differences in the sRNA content and likely the virulence factor content of Tobi-OMVs compared to ctrl-OMVs, we cannot rule out the possibility that other factors may contribute to the difference in the immune response of CF-HBECs and mouse lungs to Tobi-OMVs versus ctrl-OMVs. Nevertheless, since the inhibitor to 5′ tRNA-fMet halves transfected into CF-HBECs blocked the Tobi-OMVs mediated reduction in IL-8 secretion compared to ctrl-OMVs, we conclude that 5′ tRNA-fMet halves play a major role in suppressing IL-8 levels and neutrophil recruitment. Third, tRNAs are known to have post-transcriptional modifications, which affect RNA structure and RNA-protein interaction (Lorenz et al., 2017). Whether the modifications on 5′ tRNA-fMet halves modulate the anti-inflammatory effect requires additional study.

Highly effective CFTR modulator drugs have significantly improved outcomes in pwCF; however, in a few recent studies, they have been shown to have either no effect or a modest effect on the *P. aeruginosa* burden in the CF lungs (Davies and Martin, 2018; Yi et al., 2021). Thus, new approaches are needed to reduce the bacterial load in the lungs of chronically colonized pwCF. We propose that 5′ tRNA-fMet halves or similar miRNA-like molecules may be utilized as a therapeutic strategy to reduce IL-8 and neutrophil content in the lungs of pwCF, resulting in reduced lung damage and improved lung function.

## Methods

### *P. aeruginosa* cultures

*P. aeruginosa* (strain PA14) and clinical isolates were grown in lysogeny broth (LB, Thermo Fisher Scientific, Waltham, MA) liquid cultures at 37°C with shaking at 225 rpm. Tobramycin (1 μg/mL), a concentration that reduces *P. aeruginosa* by an amount similar to that observed clinically, or vehicle was added to the cultures. The clinical isolates, two mucoid and two non-mucoid strains, have been characterized previously (Yu et al., 2012; Moreau-Marquis et al., 2015). In some experiments, 5′ tRNA-fMet1 half (5′-CGCGGGGTGGAGCAGTCTGGTAGCTCGTCGGGCTC-3′) was cloned into the arabinose-inducible expression vector pMQ70 (Shanks et al., 2006) by cutting EcoRI and SmaI restriction sites. GenScript (GenScript USA Inc., Piscataway, NJ, USA) performed the cloning procedure. PA14 was transformed with the 5′ tRNA-fMet1 half expression vector or empty vector via electroporation. *P. aeruginosa* strains with the arabinose-inducible vector and its derivatives were grown in LB with 133 mM L-arabinose (2% w/v) and 300 μg/ml carbenicillin (both from Sigma-Aldrich).

### Growth kinetics of *P. aeruginosa*

*P. aeruginosa* overnight LB cultures were centrifuged, washed, and resuspended in fresh LB before measuring the optical density at 600 nm (OD600) to determine cell number. Bacteria were seeded at 1×10^5^ cells per 100 μl LB with or without tobramycin (1 μg/mL) in a transparent, flat bottom, 96-well plate covered with a lid. The plate was cultured in a plate reader at 37°C for 24 h. The reader was programmed to measure the OD600 every 10 minutes after shaking the plate for 5 seconds.

### Outer membrane vesicle preparation and quantification

OMVs were isolated as described by us previously (Koeppen et al., 2016; Bauman and Kuehn, 2006). Briefly, *P. aeruginosa* overnight cultures were centrifuged for 1 h at 2800 g and 4°C to pellet the bacteria. The supernatant was filtered twice through 0.45 μm PVDF membrane filters (Millipore, Billerica, MA, USA) to remove bacteria and concentrated with 30K Amicon filters (Millipore, Billerica, MA, USA) at 2800 g and 4°C to obtain ∼200 μL concentrate. The concentrate was resuspended in OMV buffer (20 mM HEPES, 500 mM NaCl, pH 7.4) and subjected to ultracentrifugation for 2 h at 200,000 g and 4°C to pellet OMVs. OMV pellets were re-suspended in 60% OptiPrep Density Gradient Medium (Sigma-Aldrich, Cat. # D1556) and layered with 40%, 35%, 30% and 20% OptiPrep diluted in OMV buffer. OMVs in OptiPrep layers were centrifuged for 16 h at 100,000 g and 4°C. 500 μl fractions were taken from the top of the gradient, with OMVs residing in fractions 2 and 3, corresponding to 25% OptiPrep. The purified OMVs were quantified by nanoparticle tracking analysis (NTA, NanoSight NS300, Malvern Panalytical Ltd, Malvern, UK) before exposure of CF-HBECs or mice to OMVs.

### CF-HBEC culture

De-identified primary human bronchial epithelial cells from four CF donors (CF-HBECs, Phe508del homozygous) were obtained from Dr. Scott Randell (University of North Carolina, Chapel Hill, NC, USA) and cultured as described previously (Fulcher and Randell, 2013; Koeppen et al., 2021). Briefly, cells were grown in BronchiaLife basal medium (Lifeline Cell Technology, Frederick, MD, USA) supplemented with the BronchiaLife B/T LifeFactors Kit (Lifeline) as well as 10,000 U/ml Penicillin and 10,000 μg/ml Streptomycin.

To polarize cells, CF-HBECs were seeded on polyester transwell permeable filters (#3405 for 24-mm transwell or #3801 for 12-mm Snapwell; Corning, Corning, NY) coated with 50 μg/ml Collagen type IV (Sigma-Aldrich, St. Louis, MO). Air Liquid Interface (ALI) medium was added to both apical and basolateral sides for cell growth. Once a confluent monolayer was obtained, the apical medium was removed, and cells were cultured at an air-liquid interface and fed basolaterally every other day with ALI media for 3-4 weeks before cells were fully polarized for treatment (Randell et al., 2011).

### Exposure of cells to OMVs

Polarized cells on 12-mm Snapwell filters were washed with PBS to remove excess mucus, and 2 mL of serum-free ALI medium was added to the basolateral side. 1.5×10^10^ purified OMVs or the same volume of Optiprep vehicle control in 200 μL serum-free ALI medium were applied to the apical side of cells. 2.1×10^10^ Tobi-OMVs (1.4X Tobi-OMVs) were also used. After a six-hour exposure, the basolateral medium was collected for cytokine measurements.

### Cytokine measurements

Cytokine secretion from primary CF-HBECs was measured with the Human IL-8/CXCL8 DuoSet ELISA (#DY208, R&D Systems, Minneapolis, MN). Several samples were also screened with MILLIPLEX MAP Human Cytokine/Chemokine 41-Plex cytokine assay (Millipore). Cytokines in mouse BALF were analyzed with the Mouse CXCL1/KC DuoSet ELISA (#DY453, R&D Systems, Minneapolis, MN).

### RNA isolation and small RNA-seq analysis

PA14 was grown in T-broth (10 g tryptone and 5 g NaCl in 1 L H_2_O) with or without tobramycin (1 μg/mL) to reduce small RNA reads from yeast present in LB medium. The culture supernatants were processed as mentioned above to obtain OMV pellets. The pellets were resuspended with OMV buffer and re-pelleted again by centrifugation at 200,000 g for 2 h at 4°C and lysed with Qiazol followed by RNA isolation with the miRNeasy kit (Qiagen) to obtain total RNA including the small RNA fraction. DNase-treated total RNA was used to prepare cDNA libraries with the SMARTer smRNA-Seq Kit (Takara Bio, Mountain View, CA). Libraries were sequenced as 50 bp single-end reads on an Illumina HiSeq sequencer. The first three nucleotides of all reads and the adapter sequences were trimmed using cutadapt (Martin, 2011) before sequence alignment.

To verify the overexpression of 5′ tRNA-fMet1 half, PA14 clones with the 5′ tRNA-fMet1 half expression plasmid or the empty pMQ70 vector were grown in LB (with L-arabinose and carbenicillin) for isolation of V-OMVs and tRNA1-OMVs. The OMV pellets were collected and processed as described above to isolate RNA. The QIAseq miRNA Library Kit (Qiagen) was used to prepare cDNA libraries, and 50 bp single-end sequencing was performed on an Illumina MiniSeq system.

Reads were aligned to the PA14 reference genome using CLC Genomics Workbench (CLC-Bio/Qiagen) with the following modifications from the standard parameters: a) the maximum number of mismatches = zero to eliminate unspecific alignment and b) the maximum number of hits for a read = 30 to capture all sRNAs aligned to the PA14 genome. Pileups of mapped reads and frequency tables for each unique sequence were exported for normalization and further analysis with the software package edgeR in the R environment (R Core Team, 2021; Robinson et al., 2010).

### Detection of 5′ tRNA-fMet halves by RT-PCR

The induction of 5′ tRNA-fMet halves by tobramycin in OMVs of different *P. aeruginosa* strains was detected by custom Taqman Small RNA Assay (#4398987, Thermo Fisher Scientific). According to the manufacturer′ s instructions, cDNA was synthesized with the TaqMan MicroRNA Reverse Transcription Kit (#4366596, Thermo Fisher Scientific). PCR amplification and detection of 5′ tRNA-fMet halves were performed using the TaqMan Universal PCR Master Mix (#4304437, Thermo Fisher Scientific) as well as custom primers and probe design to target both 5′ tRNA-fMet halves specifically.

### 5′ tRNA-fMet1 half target prediction

The miRanda microRNA target scanning algorithm (v3.3a) was used to predict human target genes of 5′ tRNA-fMet1 half (Enright et al., 2003). The 5′ tRNA-fMet1 half sequence was scanned against human RNA sequences (annotations from GRCh38.p13 assembly) with a mimimum miRanda alignment score of 150 to generate a list of predicted target genes and the corresponding interaction minimum free energies. To account for the effect of gene expression on target prediction, for each predicted target the minimum free energy was multiplied by the gene expression level (log2CPM) in polarized HBECs identified in our previous publication (Goodale et al., 2017) to obtain an energy-expression score. 1518 genes (8.4% of all human genes) with energy-expression scores small than -200 were defined as predicted targets for the Ingenuity Pathway Analysis (Krämer et al., 2014).

### Transfection of CF-HBECs with 5′ tRNA-fMet1 half and chimeric eCLIP analysis

CF-HBECs were seeded on 15 cm dishes coated with PureCol Bovine Collagen Solution (Advanced BioMatrix, Carlsbad, CA, USA) at 2.7×10^6^ cells per dish. Three days after seeding (at 80% confluence), cells were washed and fed with complete Lifeline medium with antibiotics and transfected with 100 nM 5′ tRNA-fMet1 half (#10620310, Invitrogen custom siRNA, Thermo Fisher Scientific) or 100 nM AllStars Negative Control siRNA (siNC) using HiPerFect transfection reagent (both from Qiagen). One day after transfection, cells were washed and covered with room temperature PBS before UV irradiation (254 nm, 400 mJ/cm^2^). The irradiated cells were partially digested with pre-warmed 37°C trypsin/EDTA followed by addition of cold soybean trypsin inhibitor solution to round up cells before collection with scrapers and centrifugation at 600 x g for 10 minutes at 4°C. Cell pellets were flash-frozen in liquid nitrogen and shipped to Eclipse BioInnovations for chimeric eCLIP (Eclipse BioInnovations, San Diego, CA).

The chimeric eCLIP experiment and initial data analysis were conducted by Eclipse BioInnovations (Eclipse BioInnovations, San Diego, CA) as previously described (Van Nostrand et al., 2016) with an additional ligation step to form chimeric RNA-RNA species before 3′ RNA adapter ligation. In brief, cells were lysed and digested with RNase I. For each cell pellet, an input and an immunoprecipitated sample using an anti-AGO2 antibody (Eclipse BioInnovations, San Diego, CA) were generated for cDNA library preparation followed by paired-end 150 bp sequencing on a NovaSeq platform. Non-chimeric reads were mapped to the human genome (UCSC version GRCh38/hg38), AGO2 binding clusters were identified by CLIPper (Lovci et al., 2013) in immunoprecipitated (IP) samples and normalized against the paired input sample to define significant peaks (log2 fold change ≥ 3 of normalized reads and *P* value < 0.001 determined by Fisher′s exact test). 5′ tRNA-fMet1 half-containing chimeric reads with at least 18 nt subsequences of 5′ tRNA-fMet1 half were identified, and the subsequences were trimmed before mapping to the human genome.

For targeted chimeric eCLIP, a target-specific primer 5′ GGGTGGAGCAGTCTGGTA and a sequencing adapter-specific primer were used to enrich 5′ tRNA-fMet1 half-containing cDNA from the IP sample libraries before paired-end 150 bp sequencing on a NovaSeq platform. The primer sequence was trimmed from the 5′ ends of reads, and the remainder of reads were analyzed as non-chimeric reads as described above. The significant peak regions were identified using the same cutoffs, and HOMER′ s findMotifsGenome.pl program was used for motif enrichment analysis (Heinz et al., 2010). The resulting list of target genes with significant peaks in the transcripts was used as input for Ingenuity Pathway Analysis.

### Proteomic analysis

Primary CF-HBECs were polarized on 24 mm transwell filters and washed with PBS before treatment. 2 mL serum-free ALI medium was added to the basolateral side. 2.8×10^10^ purified V-OMVs or tRNA1-OMVs in 800 μL serum-free ALI medium were applied to the apical side of cells. After a six-hour exposure, the cells were washed with PBS and detached from the transwells with pre-warmed 37°C trypsin/EDTA. Cells were pelleted and flash-frozen in liquid nitrogen for proteomic analysis.

The cell pellets were lysed in 8M urea/50mM Tris pH 8.1/100mM NaCl + protease inhibitors (Roche) and quantified by BCA assay (Pierce), followed by trypsin digestion and desalting. 40 micrograms of peptides from each pellet were labeled with unique TMT reagent isobars; the individual TMT-labeled samples were then combined and fractionated offline into 12 fractions by PFP-RP-LC (Grassetti et al., 2017), followed by analysis on a UPLC-Orbitrap Fusion Lumos tribrid instrument in SPS-MS3 mode (McAlister et al., 2014). The resulting tandem mass spectra were data-searched using Comet; TMT reporter ion intensities were summed for each protein and normalized for total intensity across all channels. Mean fold changes comparing tRNA1-OMV-exposed cells with V-OMVs-exposed cells were calculated for each protein detected in all samples. Proteins were ranked by paired t-test *P* value, and network analysis of the top 20% proteins was performed with Ingenuity Pathway Analysis.

### Transfection of CF-HBECs with 5′ tRNA-fMet halves inhibitor and OMV exposure

CF-HBECs were seeded on PureCol-coated 12-well plates (Corning Inc.) at 50,000 cells per well. Two days after seeding (∼80% confluence), cells were washed and fed with the complete Lifeline medium plus antibiotics and transfected with 50 nM custom mirVana miRNA inhibitor (inhibitor sequence: 5′-GAGCCCGACGAGCUACCAGACUGCUCCA-3′, #4464086, Thermo Fisher Scientific) or 50 nM mirVanna inhibitor negative control#1 (#4464077, Thermo Fisher Scientific) using HiPerFect transfection reagent (Qiagen). 6 hours after transfection, cells were exposed to Optiprep vehicle ctrl, ctrl-OMVs (0.4×10^10^ per well), 1.4X Tobi-OMVs (0.55×10^10^ per well) for another 6 h, and the supernatants were collected for cytokine measurements.

### Mouse exposure to OMVs

All animal experiments were approved by the Dartmouth Institutional Animal Care and Use Committee (Protocol No. 00002026). 8–9 weeks old male and female C57BL/6J mice (The Jackson Laboratory, Bar Harbor, ME, USA) were inoculated by oropharyngeal aspiration with OMVs (0.5×10^10^ OMVs per mouse) or vehicle following brief anesthesia with isoflurane. OMV concentrations were adjusted with PBS to obtain 50 μl inoculation volume. 5 h after exposure, mice were euthanized using isoflurane anesthesia, followed by cervical dislocation after breathing stops. Mice trachea were surgically exposed, and a catheter tube was inserted into the trachea and stabilized with sutures (#100–5000, Henry Schein Inc., Melville, NY, USA). The catheter was prepared by fitting a 23 gauge needle (BD #305145, Becton, Dickinson and Company, Franklin Lakes, NJ, USA) into transparent plastic tubing (BD #427411). BALF was collected by pumping 1 ml of sterile PBS into the lungs and recovered with a syringe (BD #309659). This process was repeated once to collect 2 mL of BALF.

### Human subjects and bronchoscopy

All CF subjects were enrolled in a protocol approved by the Dartmouth Hitchcock Institutional Review Board (Protocol No. 22781). CF subjects (Phe508del homozygous) prescribed with an inhaled tobramycin regimen were enrolled if they had an FEV1 > 50% predicted, and were not currently having an exacerbation. Following informed consent, local anesthesia with nebulized lidocaine was administered to the posterior pharynx. Under conscious sedation, a flexible fiberoptic bronchoscopy was performed transorally. BALF was obtained from tertiary airways. After the bronchoscopy procedure, CF subjects were monitored per institutional protocol until they were stable for discharge.

### Quantification of neutrophils in BALF

Cells in BALF samples were pelleted and resuspended in 100 μL RBC lysis buffer (Promega) for 1 min. After removing red blood cells, the total number of cells in each BALF sample was counted, and concentrations were adjusted. 2×10^5^ cells per sample were spun onto glass slides, air-dried, and stained with the Differential Quik Stain Kit (Polysciences, Warrington, PA) according to the manufacturer′ s protocol. Neutrophils were counted under 100x magnification using a microscope. The neutrophil concentration of BALF was calculated by accounting for the retrieved BALF volume and the dilution factors used to adjust the cell concentration.

### Statistics

Data were analyzed using the R software environment for statistical computing and graphics version 4.1.0 (R Core Team, 2021) and Ingenuity Pathway Analysis (Krämer et al., 2014). Statistical significance was calculated using a mixed effect linear model, Wilcoxon rank-sum tests, paired t-tests, and likelihood ratio tests on gene-wise negative binomial generalized linear models, as indicated in the figure legends. Data were visualized, and figures were created using the R package ggplot2 (Wickham, 2016, 2).

## Supplemental material

Fig. S1 contains data on the effect of Tobi-OMVs on cytokines secreted by CF-HBEC. Fig. S2 provides validation that the 5′ tRNA-fMet1 half are over-expressed in OMVs. Table S1 contains the human BALF sample collection dates.

## Data availability

Small RNA-seq and chimera eCLIP data are available from the Gene Expression Omnibus database (accession number GSE183895, GSE183897, and GSE183898).

## Author contributions

ZL, KK, AA, DAH, SAG, and BAS designed the research studies. ZL and KK conducted experiments, acquired data, and analyzed data. AA recruited human subjects, collected clinical samples and data for analysis. SAG performed the proteomic experiment. ZL prepared figures. ZL, KK, AA, DAH, SAG, and BAS wrote the manuscript. All authors contributed to the article and approved the submitted version.

## Acknowledgments

This work was supported by the Cystic Fibrosis Foundation (STANTO19G0, STANTO20PO, STANTO19R0, and HOGAN19G0), the NIH (P30-DK117469, R01HL151385, P20-GM113132, S10OD016262), and NCCC Cancer Center Core Grants (5P30 CA023108-41, P30CA023108). We thank Dr. Fred W. Kolling for advice and support on the RNA-seq experiments.

## Tables

**Supplemental Figure 1.**
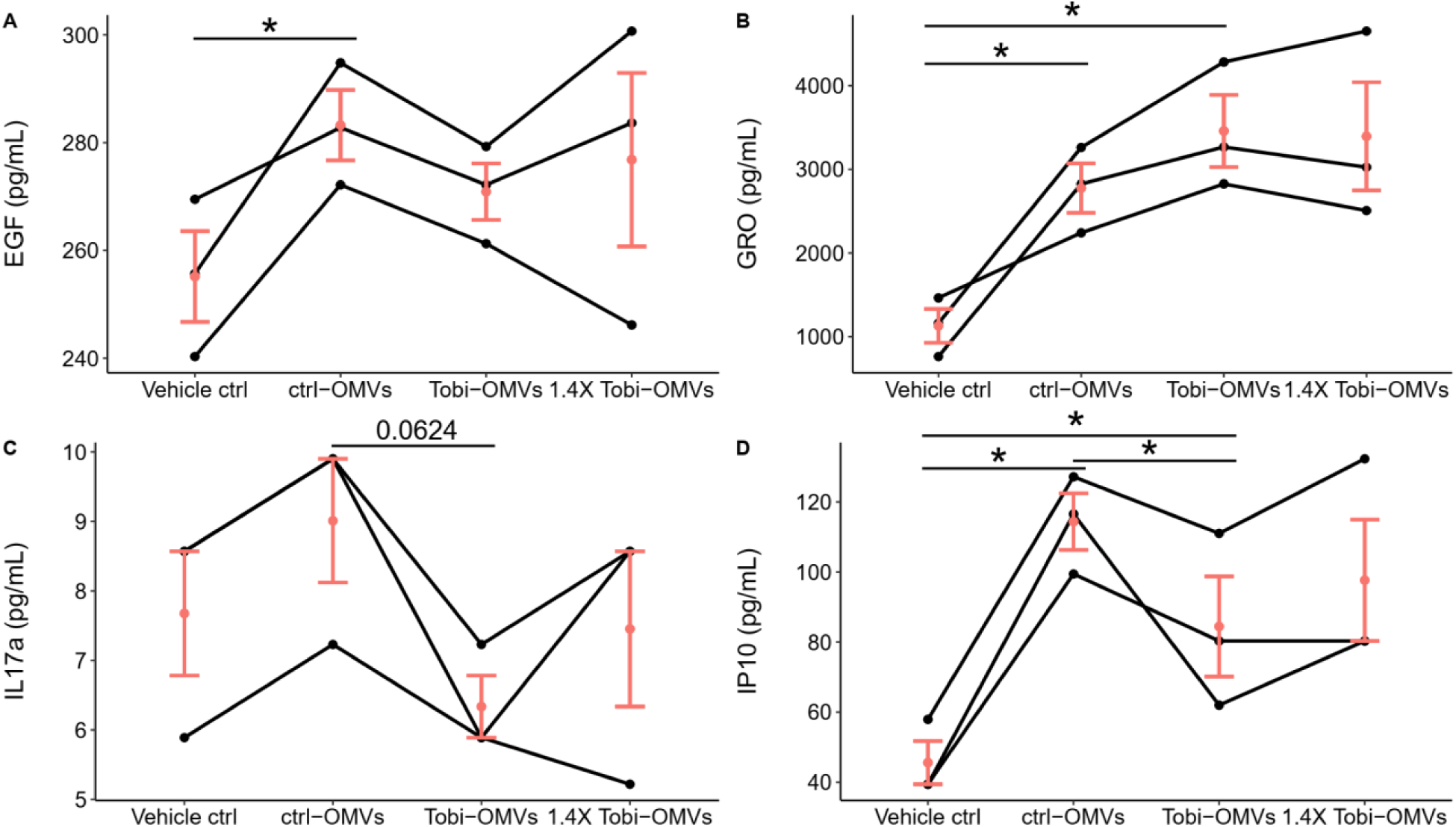
OMV effects on EGF, GRO, IL17a, and IP10. CF-HBECs from three donors (*n* = 3) were polarized in ALI culture before being exposed to either the same number of ctrl-OMVs, Tobi-OMVs, or 40% more Tobi-OMVs (1.4X Tobi-OMVs). After six-hour exposure, the basolateral medium was collected and diluted 4-fold for 41-plex measurements. Cytokines other than IL-8, EGF, GRO, IL17a, and IP10 were below the detection limit. Lines connect experiments conducted with CF-HBECs from the same donor. Horizontal red lines and red dots indicate means ± SEM. Linear mixed-effects models with CF-HBEC donor as a random effect were used to calculate *P* values; \**P* < 0.05.

**Supplemental Figure 2.**
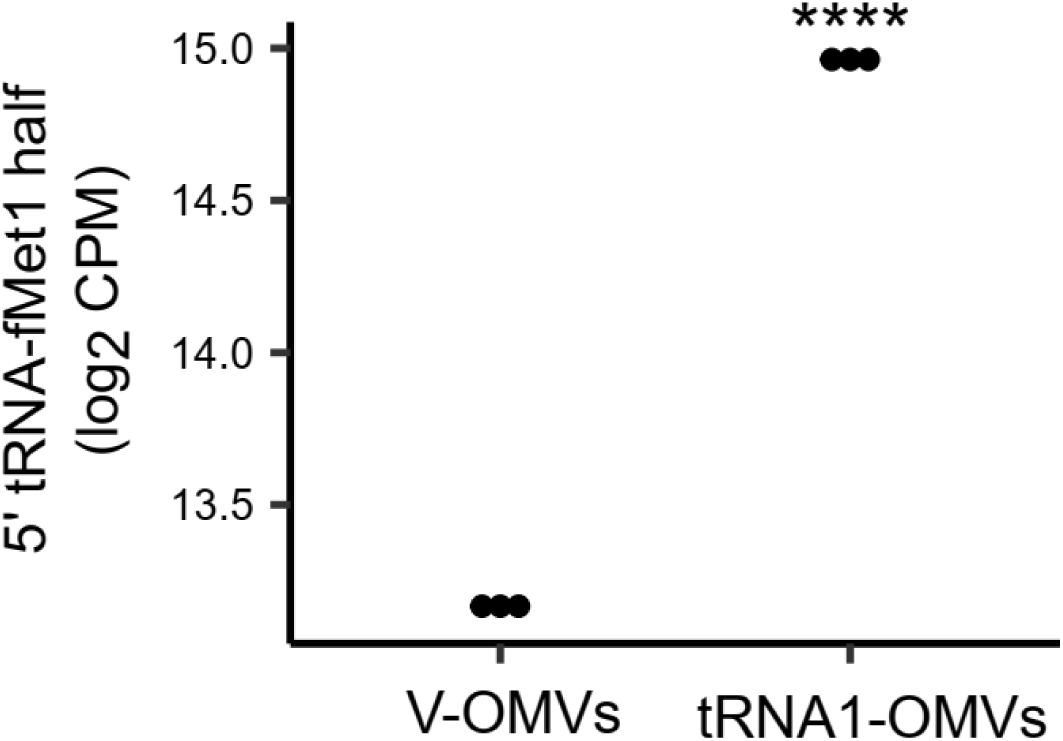
5′ tRNA-fMet1 half is over-expressed in tRNA1-OMVs compared to V-OMVs. Small RNAs in V-OMVs and tRNA1-OMVs harvested from LB culture with arabinose were subjected to small RNA sequencing to quantify 5′ tRNA-fMet1 half (*n* = 3). **** FDR < 0.0001.

**Supplemental Table 1.**
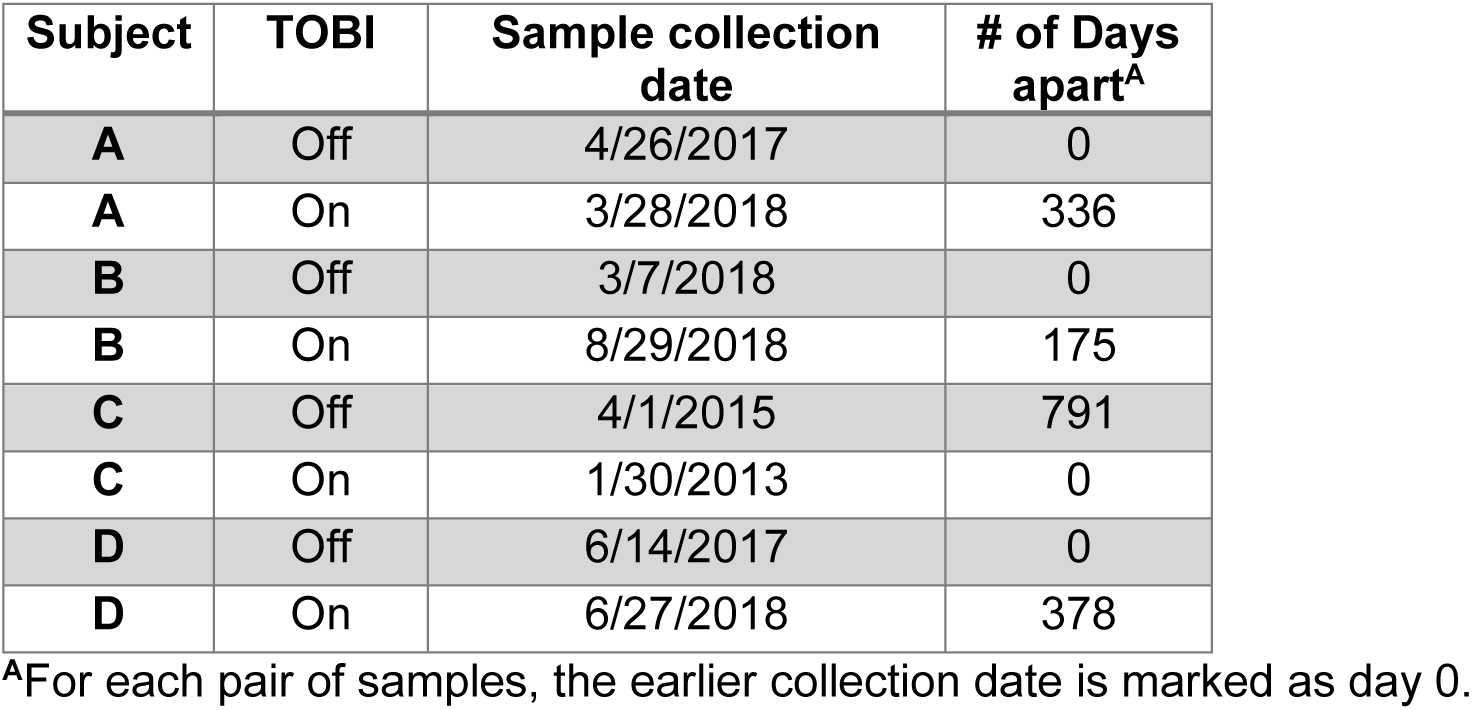
BALF sample collection dates.

| Subject | TOBI | Sample collection date | # of Days apart<sup>A</sup> |
| --- | --- | --- | --- |
| A | Off | 4/26/2017 | 0 |
| A | On | 3/28/2018 | 336 |
| B | Off | 3/7/2018 | 0 |
| B | On | 8/29/2018 | 175 |
| C | Off | 4/1/2015 | 791 |
| C | On | 1/30/2013 | 0 |
| D | Off | 6/14/2017 | 0 |
| D | On | 6/27/2018 | 378 |

<sup>A</sup>For each pair of samples, the earlier collection date is marked as day 0.

## Notes

The authors have declared that no conflict of interest exists.

### Competing Interest Statement

The authors have declared no competing interest.

